# Tumor-selective Mcl1 degradation by AUTAC uncouples antitumor efficacy from cardiotoxicity

**DOI:** 10.64898/2026.06.18.733247

**Authors:** Ahmed M. Elshazly, Janakiram R. Vangala, Adolfo G. Mauro, Fadi N. Salloum, Senthil K. Radhakrishnan

**Affiliations:** Department of Pathology, Virginia Commonwealth University, Richmond, Virginia, USA; Department of Pharmacology and Toxicology, Virginia Commonwealth University, Richmond, Virginia, USA; Department of Pharmacology and Toxicology, Faculty of Pharmacy, Kafrelsheikh University, Kafrelsheikh, Egypt; Pauley Heart Center, Department of Internal Medicine, Virginia Commonwealth University, Richmond, Virginia, USA; Massey Comprehensive Cancer Center, Virginia Commonwealth University, Richmond, Virginia, USA

**Author notes:** Correspondence: Senthil K. Radhakrishnan.

**Keywords:** AUTAC, TPD, Mcl1, Multiple myeloma (MM), Cardiotoxicity, Acute myeloid leukemia (AML)

## Abstract

Mcl1 is a major driver of therapeutic resistance across hematologic malignancies, but direct Mcl1 inhibition has been limited by on-target cardiotoxicity. Here, building on our development of an Mcl1-targeting autophagy-targeting chimera (AUTAC), we show that AUTAC-mediated degradation creates a tumor-selective therapeutic window that spares the heart. AUTAC induced robust cytotoxicity and Mcl1 degradation in multiple myeloma models, while showing minimal toxicity in cardiac cell lines, primary cardiomyocytes, and murine heart tissue. *In vivo*, AUTAC reduced tumor Mcl1 without measurably affecting cardiac Mcl1. Mechanistically, this selectivity was associated with lower expression of the p62/SQSTM1, TRAF6, and UBC13 machinery required for AUTAC activity in cardiac cells, together with lower intracellular AUTAC accumulation relative to tumor cells. AUTAC also enhanced the antitumor activity of carfilzomib and venetoclax, including in resistant models, without worsening cardiotoxicity or promoting cardiac Mcl1 loss. Compared with classical Mcl1 inhibitors, AUTAC caused markedly less cardiomyocyte death, mitochondrial depolarization, and apoptotic signaling. These findings identify AUTAC-mediated Mcl1 degradation as a cardiac-sparing strategy to target an otherwise clinically constrained vulnerability and support tumor-selective lysosomal degradation as a path to safer Mcl1-directed therapy.

## INTRODUCTION

Myeloid cell leukemia 1 (Mcl1) is a central anti-apoptotic member of the Bcl2 family whose sustained overexpression enables tumor cells to evade apoptosis and contributes to therapeutic refractoriness across multiple malignancies (Bolomsky et al., 2020; Hermanson et al., 2013; Mohiuddin, 2025; Siu et al., 2018; Wood, 2020). In multiple myeloma, Mcl1 upregulation is a major driver of resistance to proteasome inhibitors, which remain foundational components of frontline therapy. A substantial body of preclinical and translational evidence shows that pharmacologic neutralization of Mcl1 restores apoptotic competence and resensitizes refractory myeloma cells to proteasome inhibition (Al-Odat et al., 2021; Al-Odat et al., 2024; Nencioni et al., 2005; Rapino et al., 2013; Vanderkerken et al., 2019; Zhou et al., 2011). Similarly, adaptive resistance to the selective Bcl2 antagonist venetoclax has been mechanistically linked to compensatory upregulation of Mcl1 in acute myeloid leukemia, providing a strong biological rationale for combinatorial targeting of parallel anti-apoptotic dependencies (Condoluci and Rossi, 2022; Fischer et al., 2023; Mak et al., 2025; Ramsey et al., 2018; Zhao et al., 2023). Several Mcl1 inhibitors, including S63845 (Kotschy et al., 2016), AMG 176 (Caenepeel et al., 2018), UMI-77 (Abulwerdi et al., 2014), AZD-5991 (Tron et al., 2018), A-1210477 (Wang and Hao, 2019), and ABBV-467 (Yuda et al., 2023), have shown robust preclinical activity across multiple malignancies. These mechanistic insights and encouraging preclinical data have led to multiple early-phase clinical trials evaluating Mcl1-directed therapies (Desai et al., 2024) (NCT04178902, NCT03465540, NCT04543305, NCT02675452, NCT02992483). However, despite this compelling therapeutic rationale, the clinical development of Mcl1-targeted agents has been fundamentally limited by mechanism-based, dose-limiting cardiotoxicity (Condoluci and Rossi, 2022; Desai et al., 2024; Ong et al., 2022) (NCT03672695, NCT03218683, NCT03797261).

Mcl1 is indispensable for cardiomyocyte integrity and is expressed at substantially higher levels in cardiac tissue than other Bcl2 family members (Lee et al., 2010; Thomas and Gustafsson, 2013). Canonically, Mcl1 suppresses apoptosis by sequestering BAX and BAK, thereby preventing mitochondrial outer membrane permeabilization, cytochrome c release, and activation of caspase-9 and downstream executioner caspases (Dhani et al., 2021; Sancho et al., 2022; Wang et al., 2021; Widden and Placzek, 2021). Beyond its anti-apoptotic role, Mcl1 is essential for maintaining cardiac mitochondrial architecture and bioenergetic homeostasis. Cardiomyocyte-specific deletion of Mcl1 causes rapidly progressive dilated cardiomyopathy accompanied by profound mitochondrial disorganization and impaired oxidative phosphorylation, together with mitochondrial swelling, permeability transition pore opening, accumulation of dysfunctional mitochondria, and myocyte rupture (Thomas and Gustafsson, 2013; Wang et al., 2013). Moreover, Mcl1 is required for PINK1-Parkin-mediated mitophagy and for preservation of autophagic flux, processes that are critical for mitochondrial quality control and cardiomyocyte viability. Consistent with these findings, Guo et al. (Guo et al., 2018) reported that acute Mcl1 depletion produced only modest early changes in caspase-3/7 activity, LDH release, ATP levels, and cellular impedance, whereas sustained Mcl1 loss led to collapse of mitochondrial membrane potential, ultrastructural disruption, and impaired cardiomyocyte contractility. Collectively, these studies establish Mcl1 as a central determinant of cardiomyocyte survival, mitochondrial homeostasis, and contractile function, providing a mechanistic basis for the dose-limiting cardiotoxicity associated with systemic Mcl1 inhibition.

We recently established autophagy-targeting chimeras (AUTACs) as an effective strategy to induce selective Mcl1 degradation in hematologic malignancies, including multiple myeloma (Elshazly et al., 2026) and acute myeloid leukemia. In those settings, AUTAC enhanced the activity of proteasome inhibitors and venetoclax and overcame acquired therapeutic resistance. A critical unanswered question, however, is whether this strategy can preserve the antitumor potential of Mcl1 targeting while reducing the cardiac liability that has limited conventional Mcl1 inhibitors. In the present study, we show that AUTAC differentially affects malignant and cardiac models, promotes Mcl1 degradation and cytotoxicity in tumor cells, and displays lower cardiac toxicity than conventional Mcl1 inhibitors, including in combination settings. Together, these findings establish a clinically actionable framework for exploiting Mcl1 dependency while avoiding the dose-limiting cardiotoxicity that has constrained conventional Mcl1-targeted therapies.

## RESULTS

### AUTAC elicits robust cytotoxicity in multiple myeloma cells with minimal toxicity in cardiac models

We previously showed that AUTAC promotes autophagy-dependent degradation of Mcl1 and induces robust cell death in multiple myeloma cells (Elshazly et al., 2026). Consistent with those findings, AUTAC significantly reduced the viability of U266B1 and RPMI-8226 cells, while showing no detectable toxicity in H9c2 rat cardiomyocytes and only minimal effects in AC16 human cardiomyocytes **(Fig. 1A)**. These findings were supported by direct cell death measurements, which revealed pronounced AUTAC-induced cell death in both U266B1 and RPMI-8226 cells, with no measurable cytotoxicity in either cardiac cell line **(Fig. 1B)**. In multiple myeloma cells, AUTAC also increased cleaved caspase-3, consistent with induction of apoptosis, whereas even higher concentrations of AUTAC (10 μM) failed to induce detectable caspase-3 cleavage in cardiac cells relative to controls **(Fig. 1C)**. IncuCyte live-cell imaging further showed that AUTAC did not alter the growth kinetics of either cardiac cell line but markedly suppressed proliferation of U266B1 cells **(Fig. 1D)**, consistent with the increased cell death observed in Fig. 1B. To further evaluate cardiac safety, we treated primary cardiomyocytes with increasing concentrations of AUTAC and observed no evidence of cell death after either 24 or 48 h of treatment **(Fig. 1E)**.

**Figure 1.**
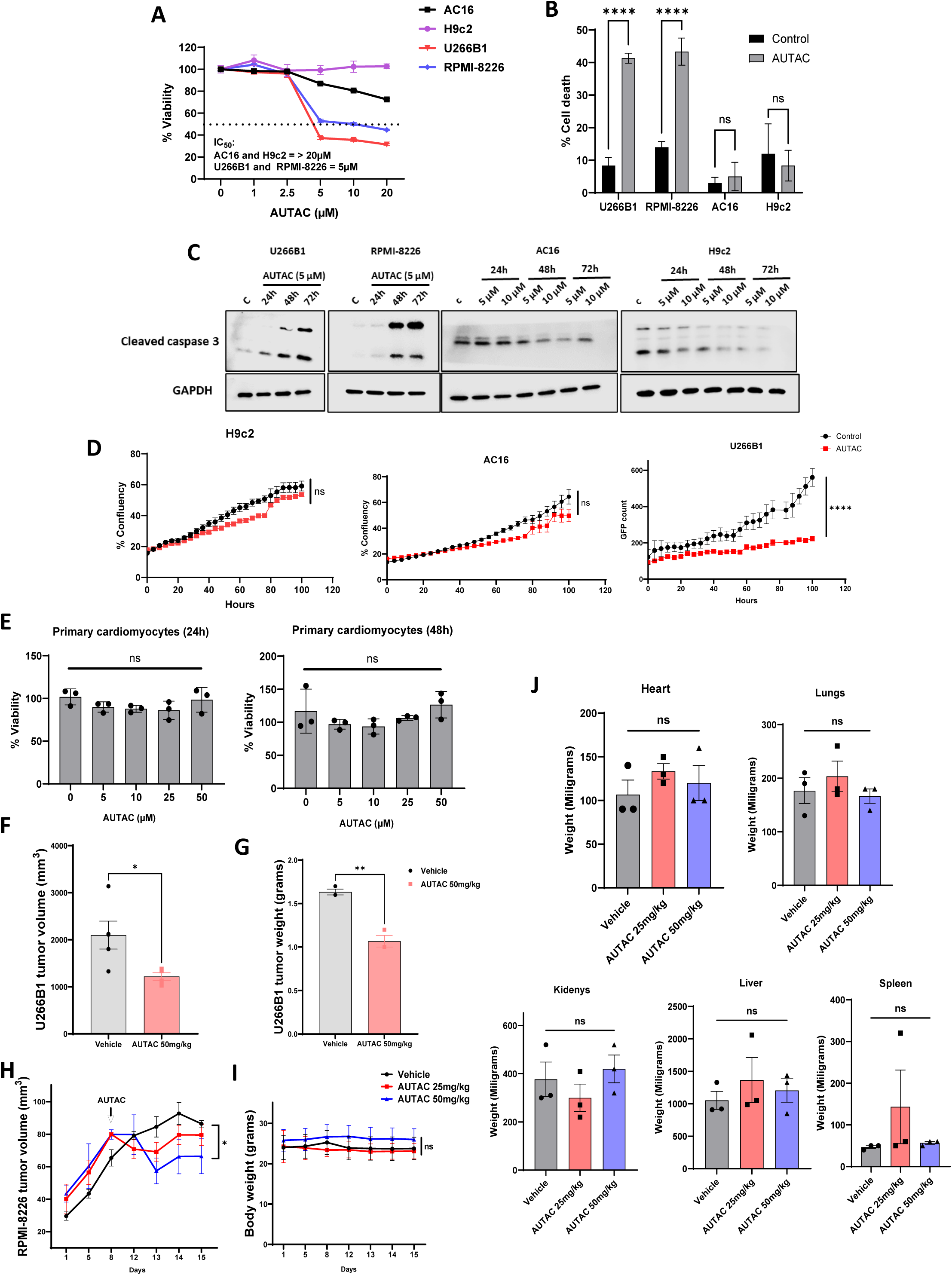
AUTAC exhibits selective cytotoxicity toward multiple myeloma cells while preserving cardiac cell viability. (A) The percentage of cell viability of U266B1, RPMI-8226, H9c2, and AC16 cells was measured after treatment with the indicated concentrations of AUTAC for 72 h. (B) The percentage of cell death of U266B1, RPMI-8226, H9c2, and AC16 cells was assessed after treatment with AUTAC (5 μM) for 72 h. (C) Western blot analysis of cleaved caspase-3 was performed in U266B1, RPMI-8226, H9c2, and AC16 cells after treatment with the indicated concentrations of AUTAC for 24, 48, and 72 h. (D) Real-time growth curves of H9c2, AC16, and U266B1 cells were monitored using the IncuCyte system. (E) The percentage of cell viability of primary cardiomyocytes was evaluated after treatment with the indicated concentrations of AUTAC for 24 and 48 h. (F) U266B1 tumor volumes from the previously reported AUTAC xenograft study (Elshazly et al., 2026); values shown correspond to the day 13 study endpoint. (G) U266B1 tumor weights from the previously reported AUTAC xenograft study (Elshazly et al., 2026); values shown correspond to the day 13 study endpoint. (H) Tumor volumes of the RPMI-8226 xenograft cohort generated for the present study after treatment with AUTAC (25 or 50 mg/kg) for 7 days. (I) Body weights of mice from the RPMI-8226 xenograft cohort shown in panel H. (J) Weights of hearts, kidneys, liver, lungs, and spleen from the RPMI-8226 xenograft cohort shown in panels H and I after treatment with AUTAC (25 or 50 mg/kg) for 8 days. Cell viability assays and Western blot experiments were conducted with three independent biological replicates per cell line. Statistical significance between experimental conditions and the indicated control or treatment groups was evaluated using an unpaired Student’s *t*-test or two-way ANOVA followed by Tukey or Sidak post hoc tests, as appropriate. Data are presented as mean ± SEM. Statistical significance is denoted as follows: \**p* < 0.05, \*\**p* < 0.01, \*\*\**p* < 0.001, and \*\*\*\**p* < 0.0001.

In the previously reported U266B1 xenograft study (Elshazly et al., 2026), analysis of the day 13 study endpoint showed that AUTAC significantly reduced tumor volume and tumor weight **(Fig. 1F and G)**. Extending these findings, in an independent RPMI-8226 xenograft cohort generated for the present study, daily administration of AUTAC (25 or 50 mg/kg) for 7 days significantly reduced tumor volume without affecting body weight **(Fig. 1H and I)**. In the same cohort, AUTAC also did not alter the weights of major organs, including the heart, kidneys, liver, lungs, and spleen **(Fig. 1J)**. Together, these data show that AUTAC exerts potent antitumor activity in multiple myeloma models while exhibiting minimal toxicity in cardiac cells and no overt systemic toxicity under the conditions tested.

### AUTAC selectively induces Mcl1 degradation in tumor cells while sparing cardiac cells

We next investigated the basis for the tumor-selective activity of AUTAC by examining its effects on Mcl1 abundance in malignant and cardiac cells. In U266B1 multiple myeloma cells, AUTAC (5 µM) induced robust Mcl1 depletion at all time points examined **(Fig. 2A)**. In contrast, AUTAC failed to alter Mcl1 protein levels in AC16 human cardiomyocytes and H9c2 rat cardiomyocytes, even at an increased concentration of 10 µM **(Fig. 2A)**. These *in vitro* findings were recapitulated *in vivo* in RPMI-8226 xenograft-bearing mice treated with AUTAC at 25 or 50 mg/kg. In tumor tissue, AUTAC reduced Mcl1 in a dose-dependent manner, with significant depletion observed at 50 mg/kg, whereas 25 mg/kg was insufficient to induce detectable Mcl1 loss **(Fig. 2B)**. In contrast, Mcl1 protein levels in cardiac tissue remained unchanged at both doses **(Fig. 2C)**, indicating that AUTAC selectively depletes tumor-associated Mcl1 *in vivo* while sparing cardiac Mcl1.

**Figure 2.**
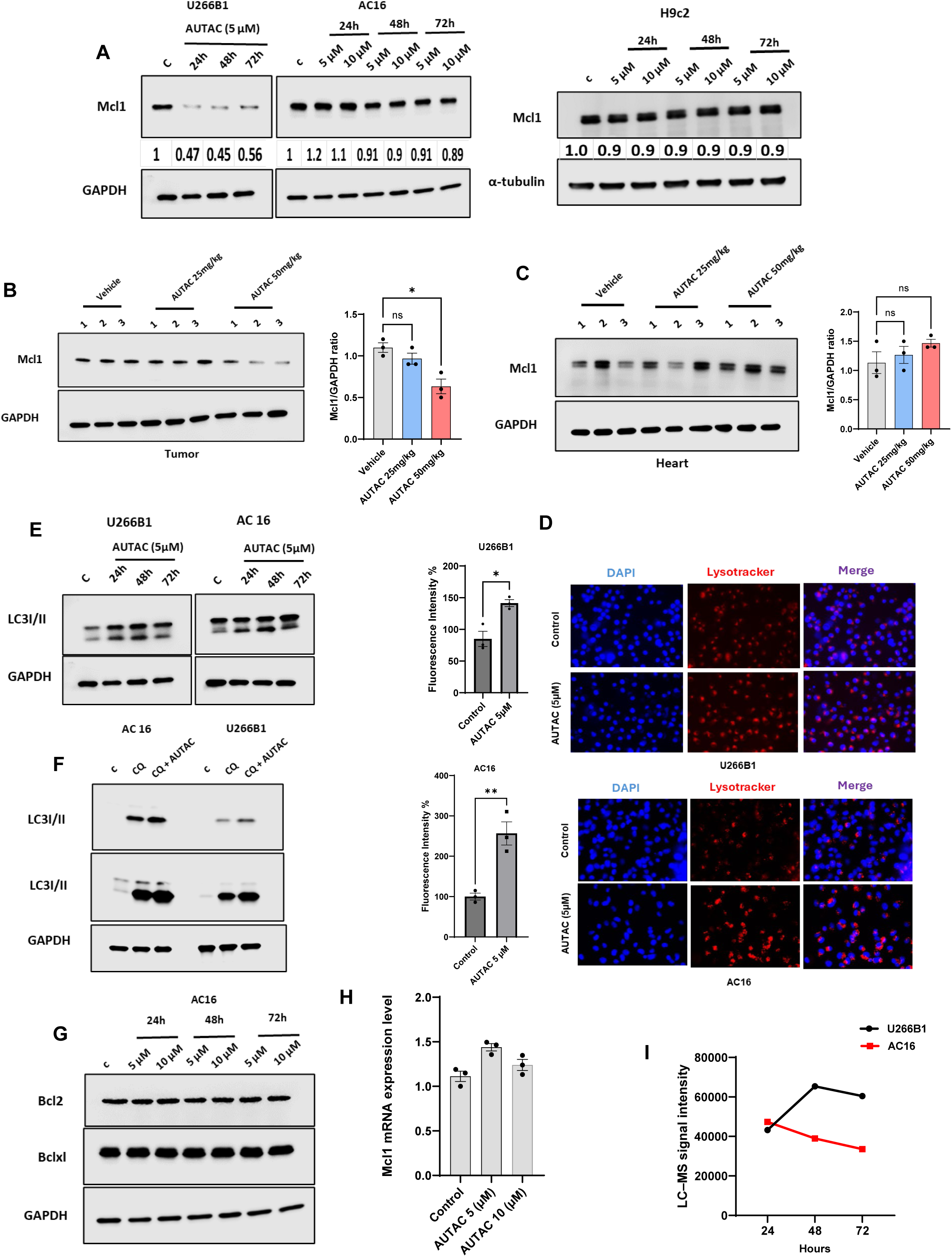
AUTAC promotes selective degradation of Mcl1 in tumor cells but not in cardiac cells. (A) Western blot analysis of Mcl1 was performed in U266B1, H9c2 and AC16 cells after treatment with the indicated concentrations of AUTAC for 24, 48, and 72 h. (B) Western blot analysis of Mcl1 was performed in RPMI-8226 tumor xenografts isolated from NSG mice treated with AUTAC (25 or 50 mg/kg) for 8 days. (C) Western blot analysis of Mcl1 was performed in hearts isolated from NSG mice treated with AUTAC (25 or 50 mg/kg) for 8 days. (D) Representative fluorescence microscopy images and quantification of U266B1 and AC16 cells treated with AUTAC (5 μM) for 72 h, followed by staining with DAPI and LysoTracker; all images were acquired under the same magnification (20×). (E) Western blot analysis of LC3I/II was performed in U266B1 and AC16 cells after treatment with AUTAC (5 μM) for 24, 48, and 72 h. (F) Western blot analysis of LC3I/II was performed in U266B1 and AC16 cells after treatment with CQ (20 μM) and/or AUTAC (5 μM) for 72 h. (G) Western blot analysis of Bcl2, and Bclxl was performed in AC16 cells after treatment with the indicated concentrations of AUTAC for 24, 48, and 72 h. (H) qRT-PCR analysis of Mcl1 mRNA expression in AC16 cells after treatment with AUTAC (5 μM or 10 μM) for 24 h. (I) LC-MS signal intensity of AUTAC in both U266B1 and AC16 cells after treatment for 72 h. Western blot experiments were conducted with three independent biological replicates per cell line. Statistical significance between experimental conditions and the indicated control or treatment groups was evaluated using an unpaired Student’s *t*-test or two-way ANOVA followed by Tukey or Sidak post hoc tests, as appropriate. Data are presented as mean ± SEM. Statistical significance is denoted as follows: \**p* < 0.05, \*\**p* < 0.01, \*\*\**p* < 0.001, and \*\*\*\**p* < 0.0001.

To define the mechanistic basis of this selectivity, we first excluded the possibility that defective autophagy in cardiomyocytes accounts for the lack of Mcl1 degradation. We previously showed that AUTAC induces Beclin1-dependent autophagy in multiple myeloma cells (Elshazly et al., 2026). Consistent with those findings, AUTAC activated autophagy in both AC16 and U266B1 cells, as evidenced by increased LysoTracker staining and accumulation of LC3I and LC3II **(Fig. 2D and E)**. To determine whether this reflected productive autophagic flux, we treated cells with chloroquine (CQ). CQ co-treatment caused greater LC3II accumulation than AUTAC alone in both tumor and cardiac cells **(Fig. 2F)**, indicating that AUTAC induces a functional and dynamic autophagic flux rather than impairing autophagosome-lysosome fusion.

We next examined whether preserved Mcl1 in cardiac cells might reflect compensatory survival signaling or transcriptional replenishment. AUTAC did not increase Bcl2 or Bclxl protein levels **(Fig. 2G)**, arguing against compensation by alternative anti-apoptotic Bcl2 family members. In addition, Mcl1 transcript levels remained unchanged following AUTAC treatment **(Fig. 2H)**, indicating that maintenance of Mcl1 protein in cardiac cells is not explained by transcriptional upregulation. We therefore asked whether differential intracellular drug accumulation might contribute to tumor selectivity. LC-MS/MS analysis confirmed intracellular accumulation of AUTAC in both U266B1 cells and AC16 cardiomyocytes **(Fig. 2I)**. Notably, AUTAC levels were approximately two-fold higher in tumor cells, particularly at 48 and 72 h after treatment, suggesting greater intracellular uptake and/or retention in the malignant compartment **(Fig. 2I)**. Together, these results show that AUTAC selectively depletes Mcl1 in tumor cells despite intact autophagic competence in cardiomyocytes and suggest that differential intracellular drug accumulation contributes to this selectivity.

### Multiple myeloma cells express higher levels of TRAF6, UBC13, and p62/SQSTM1 than cardiac cells

We previously showed that AUTAC promotes K63-linked ubiquitination of Mcl1 through the E3 ligase TRAF6 and the E2 conjugating enzyme UBC13, enabling recognition by the selective autophagy adaptor p62/SQSTM1 and delivery to LC3II-positive autophagosomes for degradation (Elshazly et al., 2026). Importantly, genetic silencing of TRAF6, UBC13, or p62/SQSTM1 abolished AUTAC-induced cytotoxicity, establishing that AUTAC functions as a degrader rather than a conventional inhibitor (Elshazly et al., 2026). To explore whether differential expression of this degradation machinery contributes to tumor selectivity, we compared the levels of these core pathway components in multiple myeloma and cardiac tissues using complementary orthogonal approaches.

Transcriptomic analysis of the MMRF CoMMpass multiple myeloma cohort and GTEx heart datasets showed significantly higher expression of p62/SQSTM1, TRAF6, and UBC13 in multiple myeloma samples than in cardiac tissue **(Fig. 3A)**. Consistent with these findings, immunoblot analysis demonstrated higher protein levels of p62/SQSTM1, TRAF6, and UBC13 in U266B1 and RPMI-8226 tumor xenografts than in cardiac tissue **(Fig. 3B and C)**. Immunohistochemical staining of U266B1 tumor xenografts and murine heart tissue further supported this pattern, showing markedly greater abundance of all three proteins in tumor tissue than in the heart **(Fig. 3D)**. A similar difference was observed at the cellular level, where U266B1 multiple myeloma cells expressed substantially higher levels of p62/SQSTM1, TRAF6, UBC13, and Mcl1 than AC16 cardiomyocytes **(Fig. 3E)**. Together, these data identify a marked enrichment of the TRAF6-UBC13-p62/SQSTM1 axis in multiple myeloma relative to cardiac tissue and support the idea that differential abundance of the AUTAC degradation machinery contributes to tumor-selective Mcl1 degradation and cytotoxicity.

**Figure 3.**
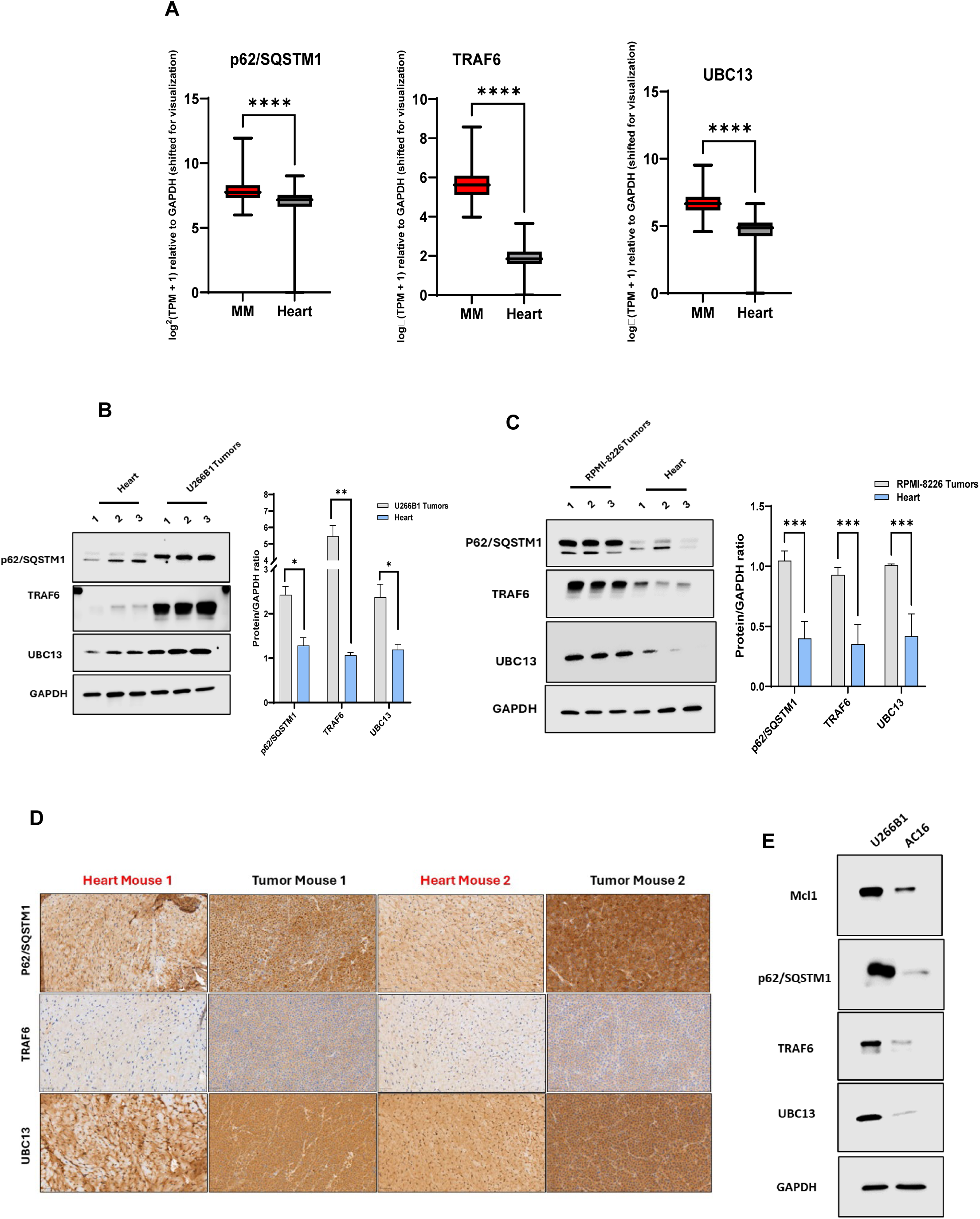
Tumor cells display elevated levels of p62/SQSTM1, TRAF6, and UBC13 relative to cardiac cells. (A) Expression levels of p62/SQSTM1, TRAF6, and UBC13 in tumor and cardiac samples from the GTEx (heart) and MMRF-CoMMpass datasets. (B) Western blot analysis of p62/SQSTM1, TRAF6, and UBC13 in U266B1 tumor xenografts and hearts isolated from NSG mice. (C) Western blot analysis of p62/SQSTM1, TRAF6, and UBC13 in RPMI-8226 tumor xenografts and hearts isolated from NSG mice. (D) Immunohistochemistry (IHC) analysis of p62/SQSTM1, TRAF6, and UBC13 in both tumor and heart tissues from NSG mice inoculated with U266B1 xenografts. (E) Western blot analysis of Mcl1, p62/SQSTM1, TRAF6, and UBC13 in U266B1 and AC16 cells. Statistical significance between experimental conditions and the indicated control or treatment groups was evaluated using an unpaired Student’s *t*-test or two-way ANOVA followed by Tukey or Sidak post hoc tests, as appropriate. Data are presented as mean ± SEM. Statistical significance is denoted as follows: \**p* < 0.05, \*\**p* < 0.01, \*\*\**p* < 0.001, and \*\*\*\**p* < 0.0001.

### AUTAC enhances the efficacy of carfilzomib and venetoclax without increasing cardiac toxicity

We next examined whether combining AUTAC with carfilzomib (CFZ) or venetoclax alters cardiac toxicity while preserving the antitumor benefit of these regimens. In cardiac cells, AUTAC did not increase CFZ-associated toxicity **(Fig. 4A)**. In contrast, AUTAC significantly enhanced the efficacy of CFZ in U266B1 multiple myeloma cells and in the proteasome inhibitor-resistant U266B1^R^ cell line **(Fig. 4B)**. These findings were supported by direct cell death measurements, in which the AUTAC-CFZ combination produced substantially greater cytotoxicity in U266B1 cells than either agent alone while sparing cardiac cells **(Fig. 4C)**. Consistent with this effect, immunoblot analysis showed marked increases in cleaved caspase-3 and cleaved PARP in U266B1 cells following combination treatment **(Fig. 4D)**, indicating enhanced apoptotic signaling. In contrast, AUTAC did not increase cleaved PARP or cleaved caspase-3 beyond the levels observed with CFZ alone in cardiac cells **(Fig. 4D)**, suggesting that AUTAC does not potentiate CFZ-associated cardiotoxic signaling.

**Figure 4.**
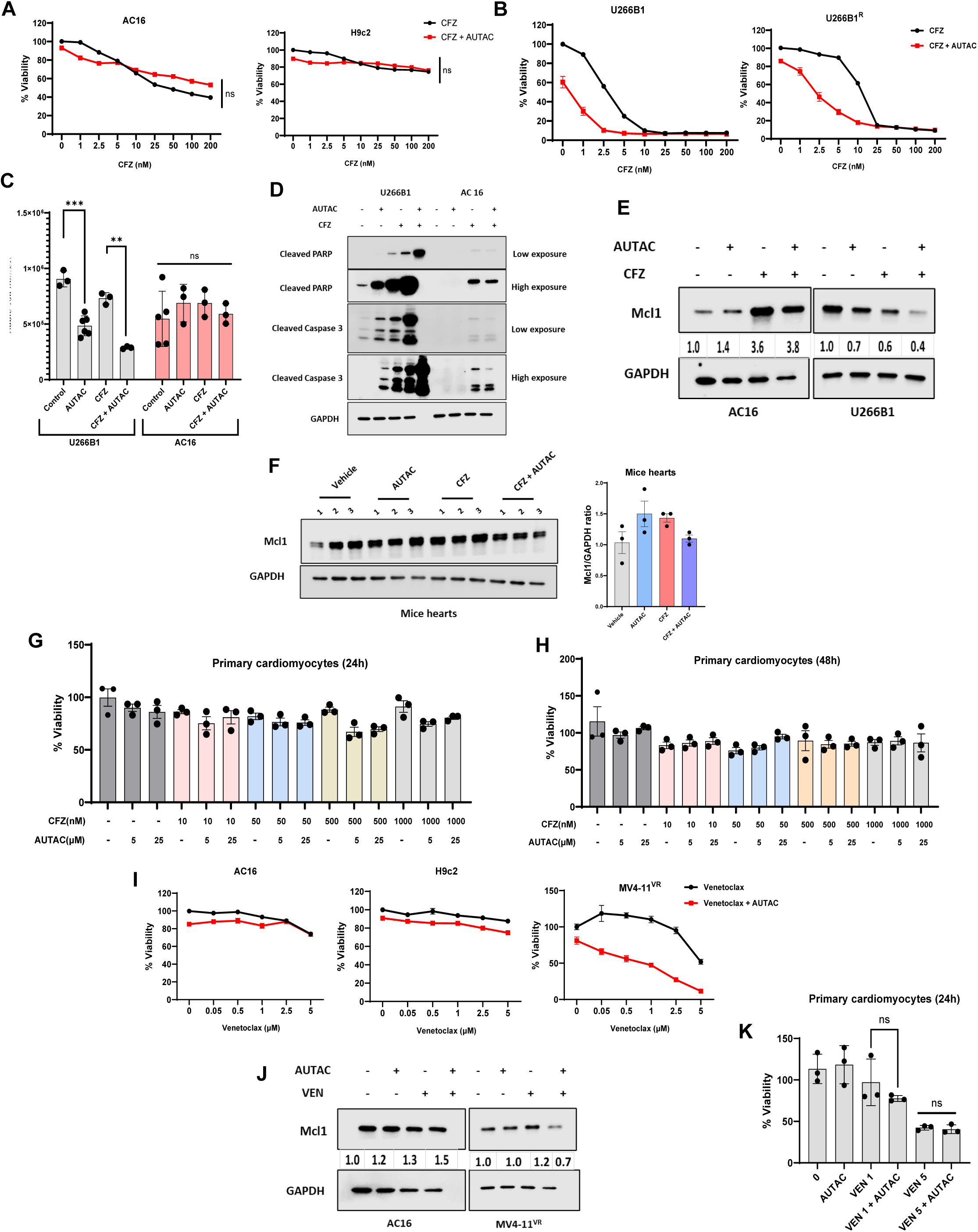
AUTAC enhances the antitumor activity of carfilzomib and venetoclax without increasing cardiac toxicity. (A) The percentage of cell viability of H9c2 and AC16 cells, and (B) U266B1 and U266B1R cells was measured after treatment with AUTAC (5 μM) for 72 h, with the indicated concentrations of CFZ added during the final 24 h. (C) The percentage of cell death of U266B1 and AC16 cells was assessed after treatment with AUTAC (5 μM) for 72 h, with CFZ (2.5 nM) added during the final 24 h. (D) Western blot analysis of cleaved caspase-3 was performed in U266B1 and AC16 cells after treatment with AUTAC (5 μM) and/or CFZ (10 nM) for 16 h. (E) Western blot analysis of Mcl1 was performed in U266B1 and AC16 cells after treatment with AUTAC (5 μM) and/or CFZ (10 nM) for 16 h. (F) Western blot analysis of cardiac Mcl1 in hearts isolated from NSG mice bearing U266B1 xenografts and treated with vehicle, CFZ (5 mg/kg), and/or AUTAC (50 mg/kg). (G and H) The percentage of cell viability of primary cardiomyocytes was evaluated after treatment with the indicated concentrations of AUTAC for 24 (G) and 48 h (H), with CFZ added during the final 24 h. (I) The percentage of cell viability of H9c2, AC16, and MV4-11^R^ cells was measured after treatment with AUTAC (5 μM) and the indicated concentrations of venetoclax for 72h. (J) Western blot analysis of Mcl1 was performed in MV4-11^R^ and AC16 cells after treatment with AUTAC (5 μM) and/or venetoclax (5 μM) for 48h. (K) The percentage of cell viability of primary cardiomyocytes was evaluated after treatment with AUTAC (5 μM) and/or the indicated concentration of venetoclax for 24h. Cell viability assays and Western blot experiments were conducted with three independent biological replicates per cell line. Statistical significance between experimental conditions and the indicated control or treatment groups was evaluated using an unpaired Student’s *t*-test or two-way ANOVA followed by Tukey or Sidak post hoc tests, as appropriate. Data are presented as mean ± SEM. Statistical significance is denoted as follows: \**p* < 0.05, \*\**p* < 0.01, \*\*\**p* < 0.001, and \*\*\*\**p* < 0.0001.

Mechanistically, co-treatment with AUTAC and CFZ produced greater Mcl1 degradation in U266B1 cells than AUTAC alone **(Fig. 4E)**. In contrast, the AUTAC-CFZ combination did not induce detectable Mcl1 degradation in AC16 cells **(Fig. 4E)**. A similar pattern was observed *in vivo,* where AUTAC combined with CFZ did not alter cardiac Mcl1 levels **(Fig. 4F)**, whereas in our previously reported U266B1 xenograft study, the same combination reduced Mcl1 in tumor tissue (Elshazly et al., 2026). These findings were further supported in primary cardiomyocytes, where co-treatment with AUTAC and CFZ did not compromise cell viability after either 24 or 48 h of exposure **(Fig. 4G and H)**. Together, these data indicate that AUTAC enhances the antitumor activity of CFZ while preserving its selective sparing of cardiac models.

We next asked whether a similar pattern was observed with venetoclax. AUTAC did not exacerbate venetoclax-associated toxicity in either AC16 or H9c2 cardiomyocyte cell lines, while significantly enhancing venetoclax efficacy in the venetoclax-resistant MV4-11^R^ cell line **(Fig. 4I)**. Mechanistically, co-treatment with AUTAC and venetoclax did not induce detectable Mcl1 degradation in cardiac cells, whereas the combination markedly promoted Mcl1 degradation in MV4-11^R^ leukemia cells **(Fig. 4J)**. These findings were further supported in primary cardiomyocytes, in which AUTAC did not increase venetoclax-induced cytotoxicity **(Fig. 4K)**. Overall, these results show that AUTAC enhances the activity of carfilzomib and venetoclax in resistant malignant models without increasing cardiac toxicity in the systems examined.

### AUTAC showed minimal cardiotoxicity compared with classical Mcl1 inhibitors

We next compared AUTAC with classical Mcl1 inhibitors, including S63845, AMG 176, AZD-5991, UMI-77, and A-1210477. Based on IC₅₀ values reported by Elshazly et al.(Elshazly et al., 2026), AUTAC, together with UMI-77 and A-1210477, exhibited the lowest potency against the U266B1 and RPMI-8226 multiple myeloma cell lines, requiring substantially higher concentrations to achieve comparable cytotoxic effects relative to the other agents **(Fig. 5A)**. Importantly, AUTAC demonstrated a markedly superior cardiotoxicity profile. After 72 h of treatment, classical Mcl1 inhibitors induced approximately 70% cell death in AC16 human cardiomyocyte cells, whereas AUTAC resulted in only 10-20% cytotoxicity **(Fig. 5B)**. Consistently, AUTAC did not disrupt mitochondrial membrane potential in AC16 cells, in contrast to all other tested Mcl1 inhibitors, which caused a pronounced loss of mitochondrial potential **(Fig. 5C)**. These findings were further corroborated by immunoblot analysis, showing that AUTAC failed to induce the accumulation of cleaved PARP or cleaved caspase-3, while treatment with classical Mcl1 inhibitors led to robust activation of these apoptotic markers in AC16 cells **(Fig. 5D)**.

**Figure 5.**
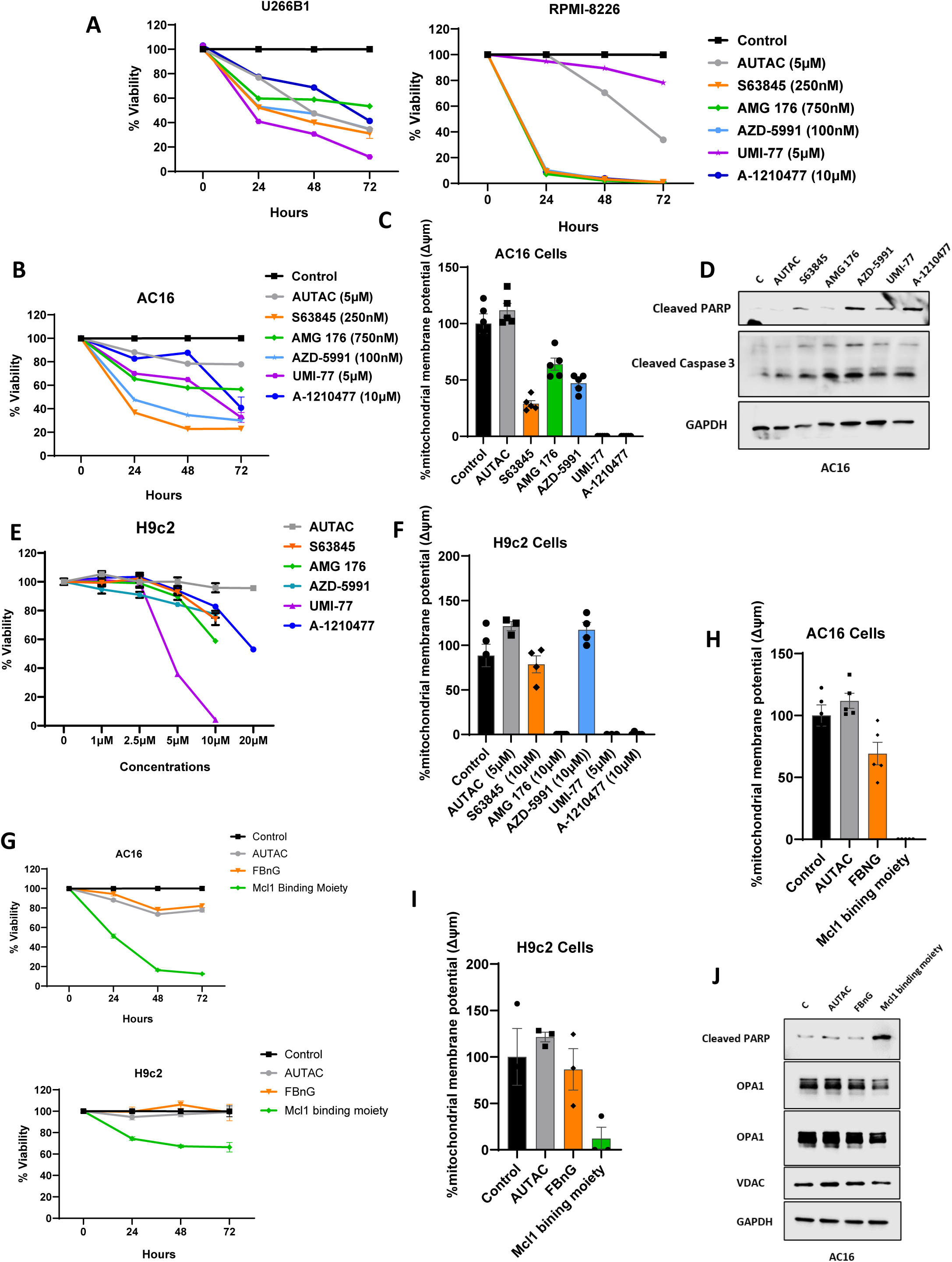
AUTAC exhibits minimal cardiotoxicity compared with traditional Mcl1 inhibitors. (A) The percentage of cell viability of U266B1 and RPMI-8226 cells was measured after treatment with the IC₅₀ values of AUTAC, S63845, AMG 176, AZD-5991, UMI-77, and A-1210477 for 24, 48, and 72 h. (B) The percentage of cell viability of AC16 cells was measured after treatment with the IC₅₀ values of AUTAC, S63845, AMG 176, AZD-5991, UMI-77, and A-1210477 for 24, 48, and 72 h. (C) TMRE analysis was performed in AC16 cells after treatment with the IC₅₀ values of the indicated compounds for 72 h. (D) Western blot analysis of cleaved caspase-3 was performed in AC16 cells after treatment with the IC₅₀ values of the indicated compounds for 48 h. (E) The percentage of cell viability of H9c2 cells was measured after treatment with the indicated concentrations of AUTAC, S63845, AMG 176, AZD-5991, UMI-77, and A-1210477 for 72 h. (F) TMRE analysis was performed in H9c2 cells after treatment with the indicated compounds for 72 h. (G) The percentage of cell viability of AC16 and H9c2 cells was measured after treatment with AUTAC (5 μM), FBnG (2.65μM), and the Mcl1 moiety (2.76 μM) for 24, 48, and 72 h. The concentrations of FBnG and the Mcl1 moiety were calculated based on their molecular weights to be equivalent to the AUTAC concentration. (H and I) TMRE analysis was performed in (H) AC16 and (I) H9c2 cells after treatment with AUTAC, FBnG, and the Mcl1 moiety for 24, 48, and 72 h. (J) Western blot analysis of cleaved PARP, OPA1, and VDAC was performed in AC16 cells after treatment with AUTAC, FBnG, and the Mcl1 moiety for 48 h. Cell viability assays, TMRE measurements and Western blot experiments were conducted with three independent biological replicates per cell line. Statistical significance between experimental conditions and the indicated control or treatment groups was evaluated using an unpaired Student’s *t*-test or two-way ANOVA followed by Tukey or Sidak post hoc tests, as appropriate. Data are presented as mean ± SEM. Statistical significance is denoted as follows: \**p* < 0.05, \*\**p* < 0.01, \*\*\**p* < 0.001, and \*\*\*\**p* < 0.0001.

To further validate these observations, we reproduced these experiments in H9c2 cardiomyocytes. AUTAC did not compromise H9c2 cell viability even at high concentrations (up to 20 µM), whereas other agents began to exhibit detectable toxicity at 10 µM **(Fig. 5E)**. Moreover, AUTAC, together with AZD-5991, did not significantly alter mitochondrial membrane potential in H9c2 cells at the tested concentrations **(Fig. 5F)**. Collectively, these data demonstrate that AUTAC displays substantially improved tolerability relative to classical Mcl1 inhibitors, particularly with respect to cardiotoxic liability.

Finally, we compared AUTAC with its individual constituent moieties, namely the FBnG autophagy-targeting module and the Mcl1-binding warhead used in the construction of the AUTAC molecule. Notably, FBnG, similar to AUTAC, did not significantly compromise the viability of AC16 human cardiomyocytes or H9c2 cardiomyocytes, whereas the Mcl1 warhead alone induced pronounced cytotoxicity **(Fig. 5G)**. These findings indicate that conjugation of the FBnG moiety to the Mcl1-binding warhead substantially mitigates the intrinsic cardiotoxicity of the Mcl1-targeting scaffold. In line with these findings, AUTAC preserved mitochondrial membrane potential in both AC16 and H9c2 cells, with FBnG exerting a partial protective effect, whereas the Mcl1-binding warhead caused a significant dissipation of mitochondrial membrane potential **(Fig. 5H and I)**. These results were further supported by immunoblot analyses, demonstrating that FBnG and AUTAC failed to induce PARP cleavage and did not compromise mitochondrial integrity, as reflected by the preserved expression levels of OPA1 and VDAC **(Fig. 5J)**. In contrast, the Mcl1-binding warhead triggered marked cytotoxicity, evidenced by robust accumulation of cleaved PARP accompanied by a substantial reduction in OPA1 and VDAC protein levels **(Fig. 5J)**. These data highlight the potential of AUTAC technology to convert intrinsically cardiotoxic (and more broadly, normal tissue-toxic) inhibitors or binders into tumor-selective targeting modalities.

## DISCUSSION

Mcl1 targeting represents one of the most compelling strategies to prevent and overcome therapeutic resistance across multiple antineoplastic modalities (Bolomsky et al., 2020; Fischer et al., 2023; Hermanson et al., 2013; Kotschy et al., 2016; Mohiuddin, 2025; Sancho et al., 2022; Siu et al., 2018; Wood, 2020). Nonetheless, the clinical promise of this approach has been undermined by mechanism-based cardiotoxicity, resulting in the premature termination of several otherwise promising clinical trials (Condoluci and Rossi, 2022; Desai et al., 2024; Ong et al., 2022). In the present study, we demonstrate that AUTAC technology can be leveraged to selectively degrade Mcl1 within malignant cell populations, thereby overcoming resistance to proteasome inhibitors and venetoclax without compromising cardiac integrity. This tumor-selective therapeutic window is supported by concordant evidence from *in vitro* cancer cell line models, primary cardiomyocytes, and *in vivo* murine studies, both in single-agent settings and in combination regimens.

Since the introduction of autophagy-targeting chimera (AUTAC) technology by the Arimoto group(Takahashi and Arimoto, 2020; Takahashi et al., 2019; Takahashi et al., 2023), AUTACs have emerged as a versatile and promising platform for the selective degradation of diverse proteins and organelles, including mitochondria (Takahashi et al., 2019), MetAP2(Takahashi et al., 2019), Tau(Takahashi et al., 2019), GPX4(Gong et al., 2025), and Mcl1 (Elshazly et al., 2026). This technology incorporates an FBnG moiety that promotes K63-linked ubiquitination of the target protein, which is recognized by the ligand (warhead) moiety, through the coordinated action of E2 and E3 ubiquitin ligases (Elshazly et al., 2026; Takahashi and Arimoto, 2020; Takahashi et al., 2019; Takahashi et al., 2023). In the context of Mcl1, we identified TRAF6 as the cognate E3 ligase and UBC13 as the requisite E2 enzyme mediating AUTAC-induced Mcl1 ubiquitination and subsequent degradation (Elshazly et al., 2026). The resulting K63-polyubiquitinated Mcl1 is then recognized by the autophagy adaptor p62/SQSTM1 via its ubiquitin-binding domain and delivered to LC3II-positive autophagosomes through its LC3-interacting region, culminating in lysosomal degradation (Elshazly et al., 2026). Furthermore, we previously demonstrated that AUTAC elicits robust cytotoxicity in multiple myeloma cells in a manner that is mechanistically dependent on the UBC13-TRAF6-p62 axis, as genetic silencing of any component of this pathway completely abrogated AUTAC-mediated cytotoxic effects (Elshazly et al., 2026). Building on these findings, the present study demonstrates that AUTAC exhibits tumor-selective cytotoxicity and preferentially induces Mcl1 degradation in malignant cells while sparing cardiomyocytes. Mechanistically, this selectivity is attributable, in part, to the comparatively low expression levels of p62/SQSTM1, TRAF6, and UBC13 in cardiomyocytes. This differential expression was validated across multiple orthogonal approaches, including analysis of publicly available transcriptomic datasets, immunoblotting in cancer and cardiac cell lines, and comparative profiling of tumor xenografts versus murine cardiac tissue.

An additional contributor to AUTAC selectivity may arise from differential cellular pharmacokinetic partitioning. Specifically, AUTAC accumulates to approximately twofold higher intracellular concentrations in tumor cells relative to cardiomyocytes, particularly at later time points (48 and 72 h). Consequently, substantially higher concentrations of AUTAC are required to achieve intracellular levels in cardiomyocytes comparable to those observed in tumor cells, further contributing to the tumor-selective therapeutic window.

Importantly, AUTAC retained its tumor-selective therapeutic window when combined with carfilzomib or venetoclax, further supporting the translational potential of this approach in combination regimens. Mechanistically, AUTAC exhibited pronounced synergy with carfilzomib and venetoclax by exploiting therapy-induced autophagic flux to accelerate and enhance Mcl1 degradation (Elshazly et al., 2026). This mechanistic convergence translated into robust synergistic cytotoxicity, particularly in models of resistance to both agents, while preserving cardiac integrity. Notably, accumulating clinical evidence has linked proteasome inhibitor therapy to cardiotoxicity in multiple myeloma, thereby constraining their therapeutic index and limiting rational combination strategies with other agents (Georgiopoulos et al., 2023; Grandin et al., 2015; Land et al., 2015; Mendez-Lopez et al., 2025; Wu et al., 2020). In the current study, we further demonstrate that AUTAC synergizes with carfilzomib, raising the possibility that lower carfilzomib doses could retain antitumor activity and potentially reduce carfilzomib-associated cardiotoxicity. Overall, these findings highlight the potential of AUTAC technology to reprogram side-effect-associated target engagement into tumor-selective degradation by leveraging differential expression of mechanistically requisite ligases and cargo receptors between malignant and healthy tissues.

## MATERIALS AND METHODS

### Cell Culture

RPMI-8226 (CCL-155; ATCC), U266B1 resistant/ U266B1^R^ (Zhao et al., 2023), U266B1 (TIB-196; ATCC), were grown in Roswell Park Memorial Institute Medium (RPMI) (Gibco), MV4-11 resistant cells/MV4-11^R^ cells were grown in Iscove’s Modified Dulbecco’s Medium (IMEM; Gibco), while AC16 (CRL-3568; ATCC) and H9c2 (CRL-1446, ATCC) cells were grown in Dulbecco’s Modified Eagle Medium (DMEM; Gibco). All media were supplemented with 10% fetal bovine serum (FBS; Atlanta Biologicals), 1X penicillin-streptomycin (Pen/Strep; Invitrogen) and 5 µg/ml Plasmocin Prophylactic (PP; InvivoGen). All cell lines were grown at 37°C in a humidified incubator with 5% CO2.

ATCC-derived cell lines were used following supplier guidelines, whereas cell lines obtained from external laboratories were used as received. Authentication was provided by the supplier. Mycoplasma contamination was routinely monitored using the InvivoGen MycoStrip® assay and prevented by Plasmocin prophylaxis.

### Primary cardiomyocytes

Hearts from anesthetized C57BL/6 mice were excised and immediately perfused with Ca²⁺-free myocyte isolation buffer using a Langendorff apparatus. The extracellular matrix was enzymatically digested using collagenase II and protease XIV. The resulting cell suspension was centrifuged at 300 × g for 2 min to collect the cardiomyocyte pellet. The pellet was gradually reintroduced to calcium using stepwise calcium reconstitution buffers prior to plating on laminin-coated culture dishes (Thermo Fisher Scientific). Cardiomyocytes were maintained in myocyte medium consisting of MEM supplemented with non-essential amino acids (NEAA), 10% fetal bovine serum (FBS), and 1% penicillin/streptomycin. After attachment, cells were treated with increasing concentrations of AUTAC and/or venetoclax for 24 h. For carfilzomib (CFZ) combination studies, cardiomyocytes were pretreated with AUTAC for 24 or 48 h, and CFZ was added during the final 24 h of treatment.

### Western blotting

Western blot was done as mentioned previously in Elshazly et al. (Elshazly et al., 2026). Briefly, cells were seeded in six-well plates and treated as indicated. At the specified time points, cells were harvested and washed once with ice-cold phosphate-buffered saline (PBS). Cell lysates were prepared using RIPA buffer, incubated on ice for 15 min, and clarified by centrifugation at 14,000 rpm for 10 min at 4 °C. Protein concentrations were determined using the bicinchoninic acid (BCA) assay. Protein samples were mixed with 4× Laemmli sample buffer containing 2-mercaptoethanol and boiled for 7 min. Equal amounts of protein (10-15 µg) were resolved by SDS-PAGE and transferred onto Immobilon-FL PVDF membranes (IPVH00010; MilliporeSigma). Membranes were blocked with 5% blotting-grade blocker (1706404; Bio-Rad) in TBST for 1-2 h at room temperature. Membranes were incubated overnight at 4 °C with the indicated primary antibodies, followed by washing with TBST and incubation with HRP-conjugated secondary antibodies for 1 h at room temperature. The following primary antibodies and dilutions were used: LC3B (2775S; Cell Signaling Technology; 1:1,000), cleaved caspase-3 (9661L; CST; 1:500), cleaved PARP (5625S; CST; 1:1,000), SQSTM1/p62 (5114S; CST; 1:1,000), Mcl1 (5453S; CST; 1:500), BCL-2 (3498S; CST; 1:1,000), BCL-xL (2764S; CST; 1:1,000), TRAF6 (8028S and 67591S; CST; 1:1,000), UBC13 (4919S; CST; 1:1,000), and GAPDH (5174S; CST; 1:10,000). HRP-conjugated goat anti-rabbit IgG secondary antibody (1706515; Bio-Rad) was used at 1:10,000. Blots were developed using Clarity Western ECL substrate (1705060 or 1705062; Bio-Rad) for 5 min and imaged on the Li-COR Odyssey Fc Imaging System. Band intensities were quantified using ImageJ software.

### Cell viability

Cells were plated at the appropriate density in 96 well plate, left to attach, then treated with the indicated conditions. The CellTiter-Glo assay (G7570; Promega) was performed according to standard protocol and luminescence was quantified using a POLARStar Optima cell microplate reader. The figures were generated using GraphPad Prism software.

### Cell death

The percentage of dead cells were calculated using trypan blue exclusion assay (Strober, 2015). Cells were treated with the indicated conditions then stained with trypan blue, then counted. The figures were generated using GraphPad Prism software.

### IncuCyte Live-Cell Imaging

Real-time monitoring of cell proliferation and cytotoxicity was performed using the IncuCyte Live-Cell Analysis System (Sartorius). Cells were seeded in 96-well plates at optimized densities and allowed to adhere overnight. The following day, cells were treated under the indicated conditions. Plates were transferred to the IncuCyte system and imaged every 4 h for up to 100 h using phase-contrast (for AC16 and H9c2) and GFP fluorescence channels (U266B1). Cell confluence and GFP counts were quantified automatically using the IncuCyte analysis software and normalized to the initial confluence at the time of treatment.

### Gene Expression Analysis from Public Datasets

Gene expression data from the GTEx Heart, and MMRF-CoMMpass cohorts were obtained via the UCSC Xena Browser. Transcript abundance values were retrieved as log₂-transformed [log₂(TPM + 1)]. To normalize for inter-sample variability, expression values were normalized to GAPDH. For visualization purposes, a constant value (+11) was added uniformly to all normalized values to eliminate negative values without altering relative differences.

### Lysotracker imaging

AC16 and U266B1 cells were plated in 12 well plates and treated with AUTAC 5 μM for 72h. Cells stained with DAPI (Cat No: PI62249; ThermoFisher) and Lysotracker (Cat No: L7528; ThermoFisher) stains according to the manufacturer’s protocol. Fluorescence images of cells were taken using Zeiss AxioObserver Inverted Microscope. The merged images and quantification were generated using Image J software. The figures were generated using GraphPad Prism software.

### Mitochondrial Membrane Potential (TMRE Assay)

Mitochondrial membrane potential was assessed in live cells using the TMRE Mitochondrial Membrane Potential Assay Kit (ab113852; Abcam), according to the manufacturer’s instructions. Cells were seeded at optimized densities and treated as indicated. TMRE working solution was freshly prepared in complete culture medium and added to cells at 700nM, followed by incubation for 30 minutes at 37 °C protected from light. Where indicated, FCCP (20 µM) was added 10 min prior to TMRE staining as a positive control for mitochondrial depolarization. After staining, cells were gently washed with PBS. TMRE fluorescence was measured using a fluorescence plate reader (Ex/Em 549/575 nm). TMRE fluorescence intensity was quantified as a relative measure of mitochondrial membrane potential and normalized to vehicle-treated controls.

### *In vivo* and immunohistochemical staining

All animal studies were approved by the Virginia Commonwealth University (VCU) Institutional Animal Care and Use Committee (IACUC) (Protocol AD10001417) and conducted in accordance with institutional guidelines and regulations. Animal experiments are reported in compliance with the ARRIVE 2.0 guidelines. Xenograft implantation, dosing, and tumor measurements were performed by the Massey Cancer Center Mouse Core.

For the RPMI-8226 xenograft study generated for the present manuscript, male and female NOD-SCID-IL2Rγ (NSG) mice (4-6 weeks old) were inoculated subcutaneously (unilaterally, right flank) with RPMI-8226 cells (5 × 10^6^ cells; n = 3 per group) resuspended in Matrigel in a total volume of 100 µL. Tumor growth was monitored biweekly by manual caliper measurements, and tumor volumes were calculated using the modified ellipsoid formula (Euhus et al., 1986): V = (L × W²)/2, where L is the longest diameter and W is the perpendicular diameter. Once tumors reached ∼75-100 mm³, mice were randomized into treatment groups and tumor volumes were measured daily. Mice received once-daily intraperitoneal injections of vehicle (10% DMSO, 40% PEG300, 5% Tween-80, and 45% saline) or AUTAC (25 mg/kg or 50 mg/kg). Tumor volume, body weight, and terminal organ weights from this cohort are presented in Fig. 1H, I, and J.

For the U266B1 xenograft experiment, the data shown in Fig. 1F and G were derived from the same xenograft cohort described in our previously reported AUTAC study (Elshazly et al., 2026). In that study, male and female NSG mice (4-6 weeks old) were inoculated subcutaneously (unilaterally, right flank) with U266B1 cells (8 × 10^6^ cells; n = 4 per group) resuspended in Matrigel in a total volume of 100 µL. Once tumors reached approximately 75-100 mm³, mice were randomly assigned to treatment groups and, starting on day 6, received once-daily intraperitoneal injections of either vehicle (10% DMSO, 40% PEG300, 5% Tween-80, and 45% saline) or AUTAC (50 mg/kg). The prior report (Elshazly et al., 2026) described the tumor growth time course through day 10; tumors were collected at the study endpoint on day 13, when volumes reached approximately 1500 mm³, and the day 13 tumor volume and tumor weight data are presented here for the first time. No animals were excluded from this cohort or from the RPMI-8226 cohort described above.

For p62/SQSTM1, UBC13 and TRAF6 levels comparison, Male and female NOD-SCID-IL2Rγ (NSG) mice (4-6 weeks old) were inoculated subcutaneously (unilaterally, right flank) with U266B1 (11× 10^6^ cells; n = 3) resuspended in Matrigel in a total volume of 100 µL. Tumor growth was monitored biweekly by manual caliper measurements, and tumor volumes were calculated using the modified ellipsoid formula (Euhus et al., 1986): V = (L × W²)/2, where L is the longest diameter and W is the perpendicular diameter. Once tumors reached ∼75-100 mm³, tumors and hearts were collected for subsequent analyses, including Western blotting and immunohistochemistry (IHC).

The immunohistochemical staining (IHC) was performed in the VCU Tissue and Data Acquisition and Analysis Core with the Leica Bond RX autostainer using BOND heat-induced epitope retrieval solution 2, EDTA-based, pH 9, for 20 min. Antibodies were diluted in Bond Primary Antibody Diluent (Leica Biosystems AR9352) and incubated for 15 min (Ube2N/Ubc13, ab25885) or 30 min (SQSTM1/p62 (ab91526), TRAF6 (ab137452)). All antibodies were diluted at 1:500. Primary antibody incubation was followed by incubation with secondary antibodies (8 min) and HRP detection with DAB (10 min) using BOND Polymer Refine Detection (Leica Biosystems DS9800). Stained slides were then imaged on the PhenoImager HT (Akoya Biosciences).

### LC-MS/MS analysis

LC-MS/MS analysis was performed using a Shimadzu LC system comprising a CBM-20A CL communication bus module, two LC-30AD CL pumps, DGU-20A3R and DGU-20A5A degassing units, a SIL-30AC CL autosampler, and a CTO-20A CL column oven. Mass spectrometric detection was carried out on an LCMS-8060 CL triple quadrupole instrument equipped with an electrospray ionization (ESI) source (Shimadzu, Japan). Data acquisition and processing were performed using LabSolutions Insight software.

Analyses were conducted in positive ionization mode (ESI+). Mobile phase A consisted of 0.1% acetic acid in water, and mobile phase B consisted of methanol:acetonitrile (1:1, v/v). Chromatographic separation was achieved using a Thermo Scientific C18 column (2.1 × 50 mm). LHTP (20 ng/mL in methanol) was used as the internal standard.

AC16 cardiomyocytes and U266B1 multiple myeloma cells were treated with AUTAC for 24, 48, and 72 h, harvested at the indicated time points, and washed with ice-cold PBS. Samples were processed for LC-MS/MS analysis by adding 20 µL of each sample to a 96-well deep-well plate, followed by 20 µL of internal standard and 200 µL of methanol. Samples were vortexed thoroughly and centrifuged at 16,000 × g for 5 min to precipitate proteins. The clarified supernatants were transferred to autosampler vials and injected into the LC-MS/MS system for quantitative analysis.

### Quantitative Real-Time PCR

AC16 cardiomyocytes were seeded in six-well plates and grown to approximately 70% confluence at the time of treatment. Cells were exposed to AUTAC (5 or 10 µM) for 24 h. Following treatment, cells were washed once with phosphate-buffered saline (PBS) and harvested by gentle scraping in 1 mL PBS. Total RNA was isolated using the RNeasy Mini Kit (Qiagen) with on-column DNase I digestion to remove genomic DNA contamination. Complementary DNA (cDNA) was synthesized from 1 µg of total RNA using iScript Reverse Transcription Supermix (Bio-Rad), according to the manufacturer’s instructions. Quantitative real-time PCR (qRT-PCR) was performed using iTaq Universal SYBR Green Supermix (Bio-Rad) on a C1000 Touch Thermal Cycler (Bio-Rad). Transcript levels were normalized to 18S rRNA as the internal reference gene. Relative gene expression was calculated using the ΔΔCt method and analyzed with CFX Manager software (v3.1; Bio-Rad). The following human-specific primer pairs were used: Mcl1 forward, GTTTTCAGCGACGGCGTAAC, and reverse, ACTCCACAAACCCATCCCAG; 18S rRNA forward, ATGGCCGTTCTTAGTTGGTG, and reverse, CGCTGAGCCAGTCAGTGTAG.

### Statistical analysis

Unless otherwise indicated, all quantitative data is shown as mean ± SEM from at least three independent experiments, all of which were conducted in triplicates. GraphPad Prism 10 software was used for statistical analysis. Statistical significance of each condition compared to the indicated control or treatment was determined using unpaired, two-sided Student’s t-tests or two-way ANOVA with Tukey’s or Sidak’s post hoc tests, as appropriate. A P value < 0.05 was considered statistically significant. Statistical significance levels are indicated in the figures as follows: *P < 0.05; **P < 0.01; ***P < 0.001; and ****P < 0.0001.

## ACKNOWLEDGEMENTS

This work was supported by a NIH/NIGMS grant R01GM132396 (S.K.R) and an American Cancer Society Grant RSG-21-036-01-TBE (S.K.R). We thank Bin Hu, Wang Li and Vita Kraskauskiene for their assistance with the mouse studies. Services and products in support of the research project were generated by the Virginia Commonwealth University Cancer Mouse Models Core Laboratory and Tissue and Data Acquisition and Analysis Core Laboratory, supported, in part, with funding to the Massey Cancer Center from NIH-NCI Cancer Center Support Grant P30 CA016059.

## DATA AVAILABILITY STATEMENT

All data that supports the findings of this study are available from the corresponding author upon reasonable request.

## REFERENCES

Abulwerdi, F., Liao, C., Liu, M., Azmi, A.S., Aboukameel, A., Mady, A.S., Gulappa, T., Cierpicki, T., Owens, S., Zhang, T., et al. (2014). A novel small-molecule inhibitor of mcl-1 blocks pancreatic cancer growth in vitro and in vivo. Mol Cancer Ther 13, 565–575.

Al-Odat, O., von Suskil, M., Chitren, R., Elbezanti, W., Srivastava, S., Budak-Alpddogan, T., Jonnalagadda, S., Aggarwal, B., and Pandey, M. (2021). Mcl-1 Inhibition: Managing Malignancy in Multiple Myeloma. Front Pharmacol 12, 699629.

Al-Odat, O.S., Elbezanti, W.O., Gowda, K., Srivastava, S.K., Amin, S.G., Jonnalagadda, S.C., Budak-Alpdogan, T., and Pandey, M.K. (2024). KS18, a Mcl-1 inhibitor, improves the effectiveness of bortezomib and overcomes resistance in refractory multiple myeloma by triggering intrinsic apoptosis. Front Pharmacol 15, 1436786.

Bolomsky, A., Vogler, M., Köse, M.C., Heckman, C.A., Ehx, G., Ludwig, H., and Caers, J. (2020). MCL-1 inhibitors, fast-lane development of a new class of anti-cancer agents. Journal of Hematology & Oncology 13, 173.

Caenepeel, S., Brown, S.P., Belmontes, B., Moody, G., Keegan, K.S., Chui, D., Whittington, D.A., Huang, X., Poppe, L., Cheng, A.C., et al. (2018). AMG 176, a Selective MCL1 Inhibitor, Is Effective in Hematologic Cancer Models Alone and in Combination with Established Therapies. Cancer Discov 8, 1582–1597.

Condoluci, A., and Rossi, D. (2022). Mechanisms of resistance to venetoclax. Blood 140, 2094–2096.

Desai, P., Lonial, S., Cashen, A., Kamdar, M., Flinn, I., O’Brien, S., Garcia, J.S., Korde, N., Moslehi, J., Wey, M., et al. (2024). A Phase 1 First-in-Human Study of the MCL-1 Inhibitor AZD5991 in Patients with Relapsed/Refractory Hematologic Malignancies. Clin Cancer Res 30, 4844–4855.

Dhani, S., Zhao, Y., and Zhivotovsky, B. (2021). A long way to go: caspase inhibitors in clinical use. Cell Death & Disease 12, 949.

Elshazly, A.M., Hosseini, N., Vangala, J., Shen, S., Neely, V., Hu, X., Pagare, P.P., Harada, H., Grant, S., and Radhakrishnan, S.K. (2026). Proteasome inhibition enhances lysosome-mediated targeted protein degradation. Cell Death Dis.

Euhus, D.M., Hudd, C., Laregina, M.C., and Johnson, F.E. (1986). Tumor measurement in the nude mouse. Journal of surgical oncology 31, 229–234.

Fischer, M.A., Song, Y., Arrate, M.P., Gbyli, R., Villaume, M.T., Smith, B.N., Childress, M.A., Stricker, T.P., Halene, S., and Savona, M.R. (2023). Selective inhibition of MCL1 overcomes venetoclax resistance in a murine model of myelodysplastic syndromes. Haematologica 108, 522–531.

Georgiopoulos, G., Makris, N., Laina, A., Theodorakakou, F., Briasoulis, A., Trougakos, I.P., Dimopoulos, M.A., Kastritis, E., and Stamatelopoulos, K. (2023). Cardiovascular Toxicity of Proteasome Inhibitors: Underlying Mechanisms and Management Strategies: JACC: CardioOncology State-of-the-Art Review. JACC CardioOncol 5, 1–21.

Gong, R., Wan, X., Jiang, S., Guan, Y., Li, Y., Jiang, T., Chen, Z., Zhong, C., He, L., Xiang, Z., et al. (2025). GPX4-AUTAC induces ferroptosis in breast cancer by promoting the selective autophagic degradation of GPX4 mediated by TRAF6-p62. Cell Death Differ 32, 2022–2037.

Grandin, E.W., Ky, B., Cornell, R.F., Carver, J., and Lenihan, D.J. (2015). Patterns of cardiac toxicity associated with irreversible proteasome inhibition in the treatment of multiple myeloma. J Card Fail 21, 138–144.

Guo, L., Eldridge, S., Furniss, M., Mussio, J., and Davis, M. (2018). Role of Mcl-1 in regulation of cell death in human induced pluripotent stem cell-derived cardiomyocytes in vitro. Toxicology and Applied Pharmacology 360, 88–98.

Hermanson, D.L., Das, S.G., Li, Y., and Xing, C. (2013). Overexpression of Mcl-1 confers multidrug resistance, whereas topoisomerase IIβ downregulation introduces mitoxantrone-specific drug resistance in acute myeloid leukemia. Mol Pharmacol 84, 236–243.

Kotschy, A., Szlavik, Z., Murray, J., Davidson, J., Maragno, A.L., Le Toumelin-Braizat, G., Chanrion, M., Kelly, G.L., Gong, J.N., Moujalled, D.M., et al. (2016). The MCL1 inhibitor S63845 is tolerable and effective in diverse cancer models. Nature 538, 477–482.

Land, J., Afifi, S., Adel, N.G., Devlin, S., Arora, A., Lendvai, N., and Landgren, O. (2015). Incidence and Management of Proteasome Inhibitor-Related Cardiotoxicity in Multiple Myeloma Patients at Memorial Sloan Kettering Cancer Center. Blood 126, 4265–4265.

Lee, Y., Huang, C., Thomas, R.L., Quinsay, M.N., Rikka, S., Kubli, D.A., Fischer, K.M., Sussman, M.A., and Gustafsson, Å.B. (2010). Abstract 20062: The Anti-Apoptotic Protein Mcl-1 is an Important Regulator of Mitochondrial Turnover. Circulation 122, A20062–A20062.

Mak, P.Y., Maiti, A., Tomczyk, T., Cottens, S., Walczak, M., Andreeff, M., and Carter, B. (2025). Cardio-safe degrader CT-03p degrades MCL-1 and overcomes venetoclax resistance in AML. Blood 146, 3266.

Mendez-Lopez, M., Besse, A., Zuppinger, C., Perez-Shibayama, C., Gil-Cruz, C., Florea, B.I., De Martin, A., Lütge, M., Beckerova, D., Klimovic, S., et al. (2025). Carfilzomib-specific proteasome β5/β2 inhibition drives cardiotoxicity via remodeling of protein homeostasis and the renin-angiotensin-system. iScience 28.

Mohiuddin, M. (2025). Targeting MCL-1 to Overcome Therapeutic Resistance and Improve Cancer Mortality. Health Sci Rep 8, e71390.

Nencioni, A., Hua, F., Dillon, C.P., Yokoo, R., Scheiermann, C., Cardone, M.H., Barbieri, E., Rocco, I., Garuti, A., Wesselborg, S., et al. (2005). Evidence for a protective role of Mcl-1 in proteasome inhibitor-induced apoptosis. Blood 105, 3255–3262.

Ong, F., Kim, K., and Konopleva, M.Y. (2022). Venetoclax resistance: mechanistic insights and future strategies. Cancer Drug Resist 5, 380–400.

Ramsey, H.E., Fischer, M.A., Lee, T., Gorska, A.E., Arrate, M.P., Fuller, L., Boyd, K.L., Strickland, S.A., Sensintaffar, J., Hogdal, L.J., et al. (2018). A Novel MCL1 Inhibitor Combined with Venetoclax Rescues Venetoclax-Resistant Acute Myelogenous Leukemia. Cancer Discovery 8, 1566–1581.

Rapino, F., Naumann, I., and Fulda, S. (2013). Bortezomib antagonizes microtubule-interfering drug-induced apoptosis by inhibiting G2/M transition and MCL-1 degradation. Cell Death & Disease 4, e925–e925.

Sancho, M., Leiva, D., Lucendo, E., and Orzáez, M. (2022). Understanding MCL1: from cellular function and regulation to pharmacological inhibition. Febs j 289, 6209–6234.

Siu, K.T., Huang, C., Panaroni, C., Mukaihara, K., Fulzele, K., Soucy, R., Cidado, J., Drew, L., Chattopadhyay, S., and Raje, N. (2018). Overcoming MCL1 Resistance in Multiple Myeloma. Blood 132, 472–472.

Strober, W. (2015). Trypan Blue Exclusion Test of Cell Viability. Curr Protoc Immunol 111, A3.B.1–a3.B.3.

Takahashi, D., and Arimoto, H. (2020). Targeting selective autophagy by AUTAC degraders. Autophagy 16, 765–766.

Takahashi, D., Moriyama, J., Nakamura, T., Miki, E., Takahashi, E., Sato, A., Akaike, T., Itto-Nakama, K., and Arimoto, H. (2019). AUTACs: Cargo-Specific Degraders Using Selective Autophagy. Molecular Cell 76, 797–810.e710.

Takahashi, D., Ora, T., Sasaki, S., Ishii, N., Tanaka, T., Matsuda, T., Ikeda, M., Moriyama, J., Cho, N., Nara, H., et al. (2023). Second-Generation AUTACs for Targeted Autophagic Degradation. J Med Chem 66, 12342–12372.

Thomas, R.L., and Gustafsson, A.B. (2013). MCL1 is critical for mitochondrial function and autophagy in the heart. Autophagy 9, 1902–1903.

Tron, A.E., Belmonte, M.A., Adam, A., Aquila, B.M., Boise, L.H., Chiarparin, E., Cidado, J., Embrey, K.J., Gangl, E., Gibbons, F.D., et al. (2018). Discovery of Mcl-1-specific inhibitor AZD5991 and preclinical activity in multiple myeloma and acute myeloid leukemia. Nature Communications 9, 5341.

Vanderkerken, K., De Veirman, K., Maes, K., Menu, E., and De Bruyne, E. (2019). MCL1 Inhibitors in Multiple Myeloma. Blood 134, SCI-12-SCI-12.

Wang, H., Guo, M., Wei, H., and Chen, Y. (2021). Targeting MCL-1 in cancer: current status and perspectives. Journal of Hematology & Oncology 14, 67.

Wang, Q., and Hao, S. (2019). A-1210477, a selective MCL-1 inhibitor, overcomes ABT-737 resistance in AML. Oncol Lett 18, 5481–5489.

Wang, X., Bathina, M., Lynch, J., Koss, B., Calabrese, C., Frase, S., Schuetz, J.D., Rehg, J.E., and Opferman, J.T. (2013). Deletion of MCL-1 causes lethal cardiac failure and mitochondrial dysfunction. Genes Dev 27, 1351–1364.

Widden, H., and Placzek, W.J. (2021). The multiple mechanisms of MCL1 in the regulation of cell fate. Communications Biology 4, 1029.

Wood, K.C. (2020). Overcoming MCL-1-driven adaptive resistance to targeted therapies. Nature Communications 11, 531.

Wu, P., Oren, O., Gertz, M.A., and Yang, E.H. (2020). Proteasome Inhibitor-Related Cardiotoxicity: Mechanisms, Diagnosis, and Management. Curr Oncol Rep 22, 66.

Yuda, J., Will, C., Phillips, D.C., Abraham, L., Alvey, C., Avigdor, A., Buck, W., Besenhofer, L., Boghaert, E., Cheng, D., et al. (2023). Selective MCL-1 inhibitor ABBV-467 is efficacious in tumor models but is associated with cardiac troponin increases in patients. Commun Med (Lond) 3, 154.

Zhao, T., He, Q., Xie, S., Zhan, H., Jiang, C., Lin, S., Liu, F., Wang, C., Chen, G., and Zeng, H. (2023). A novel Mcl-1 inhibitor synergizes with venetoclax to induce apoptosis in cancer cells. Molecular Medicine 29, 10.

Zhou, W., Hu, J., Tang, H., Wang, D., Huang, X., He, C., and Zhu, H. (2011). Small interfering RNA targeting mcl-1 enhances proteasome inhibitor-induced apoptosis in various solid malignant tumors. BMC Cancer 11, 485.

